# Potentially zoonotic gastrointestinal nematodes co-infecting free ranging non-human primates in Kenyan urban centres

**DOI:** 10.1101/2020.08.19.254714

**Authors:** Peris Mbuthia, Edwin Murungi, Vincent Owino, Mercy Akinyi, Gillian Eastwood, Richard Nyamota, Isaac Lekolool, Maamun Jeneby

## Abstract

**Background:** Natural infections with soil transmitted nematodes occur in non-human primates (NHPs) and have the potential to cross primate-species boundaries and cause diseases of significant public health concern. Despite their presence in most urban centres in Kenya, comprehensive studies on their gastrointestinal parasites are scant.

**Objective:** Conduct a cross-sectional survey to identify zoonotic nematodes in free-ranging NHPs found within four selected urban and peri-urban centres in Kenya.

**Methods:** A total of 86 NHPs: 41 African green monkeys [AGM] (*Chlorocebus aethiops*), 30 olive baboons (*Papio anubis*), 5 blue monkeys (*Cercopithecus mitis stuhlmanni*) and 10 red tailed monkeys (*Cercopithecus ascanius*) were sampled once *in situ* and released back to their habitat. Microscopy was used to identify nematodes egg and larvae stages in the samples. Subsequently, PCR coupled with high-resolution melting (PCR-HRM) analysis and sequencing were used to identify nodule worms.

**Results:** NHPs inhabiting densely populated urban environs in Kenya were found infected with a rich diversity of nematodes including three potentially zoonotic nematodes including *Oesophagostomum stephanostomum, Oesophagostomum bifurcum* and *Trichostrongylus colubriformis* and co-infections were common.

**Conclusion:** Phylogenetic analysis showed that *O. stephanostomum* from red tailed and blue monkeys have a close evolutionary relatedness to human isolates suggesting the zoonotic potential of this parasite. Moreover, we also report the first natural co-infection of *O. bifurcum* and *O. stephanostomum* in free-ranging AGMs.

## Introduction

Free-ranging non-human primates (NHPs) are a source of information regarding maintenance, transmission and disease dynamics of public health importance (Kooriyama et al., 2012; Kouassi et al., 2015; Eastwood et al, 2017). Of concern are soil-transmitted helminths (STHs) whose partial development outside their hosts allows persistence of infective stages in the environment, enabling transmission between closely related host species (Ghai et al., 2014; Cibot et al., 2015). Several studies have described cases of zoonotic STHs transmission in sympatric populations of NHPs and humans (Cibot et al., 2015; Ghai et al., 2014; Frias et al., 2019). However, little is known about urban zoonoses and possible reservoir hosts despite having abundant wildlife in most urban habitats in Africa.

Relevant to this context are the strongylid nematodes (genus *Oesophagostomum*) commonly referred to as nodular worm that parasitise pigs, ruminants, NHPs and humans. *Oesophagostomum bifurcum* is considered as the principal nodular worm of humans (Terio et al., 2018) while *Oesophagostomum stephanostomum* infects great apes including chimpanzees (Cibot et al., 2015) and gorillas (Makouloutou et al., 2014). Although the potential for cross-transmission of *Oesophagostomum* spp between humans and NHPs has been disputed (Gruijter et al., 2005; Van Lieshout et al., 2005) a novel *Oesophagostomum* clade infecting humans and sympatric NHPs populations has been described in Uganda (Ghai et al., 2014; Cibot et al., 2015). Therefore, localised research to determine the zoonotic risk and the role of NHPs as potential reservoirs, particularly in East Africa’s urban centres is imperative.

To formulate effective control measures, accurate diagnosis and genetic characterisation of nematodes is vital (Pouillevet et al., 2017). Comparatively, the polymerase chain reaction (PCR) technique is superior to conventional microscopy. Additionally, real-time high-resolution melting (HRM) analysis, a probe-free post-PCR analysis, allows direct characterisation of PCR amplicons (Reed et al., 2007). PCR-HRM has effectively been used to genotype hookworms without the need for sequencing (Ngui et al., 2012). Thus, PCR-HRM is a robust method for answering epidemiological questions that underpin pathogen surveillance and control programs (Villinger et al., 2016).

Previous surveys on helminths have focused on NHPs within wildlife reserves and rural forest habitats (Ghai et al., 2014; Akinyi et al., 2019; Obanda et al., 2019) leaving information gap on helminth zoonoses originating from free-ranging NHPs within urban and peri-urban centres. In East Africa, the sharing of habitats between NHPs and humans, such as public parks in major cities or watering points in peri-urban towns, may facilitate the cross-species transmission of zoonotic STHs. In Kenya, frequent close contacts between humans and monkeys often occurs in most urban public parks. Resident troops of monkeys often snatch food snacks from visitors, and this act may contaminate salvaged food with nematode infective stages. In addition, visitors often encourage the monkeys to climb on their heads and shoulders for photo shoot, which may represent spread of zoonotic parasites from NHPs to humans via the faecal-oral route. This human and monkey interaction suggests an increased risk for cross-transmission of zoonotic pathogens, and potential epidemiological consequences on pathogen evolution within urban ecology. Determination of the zoonotic risk and the role of NHPs as potential parasite reservoirs in urban centres is imperative. Thus, we utilised PCR-HRM and sequencing to investigate the distribution and characterise potentially zoonotic nodular worms in free-ranging NHPs within densely populated urban and peri-urban centres in Kenya.

## Materials and methods

### Study sites

This study focused on NHPs found in urban centres of Mombasa and Kisumu, and peri-urban centres within Murang’a and Kakamega counties of Kenya (Figure 1). NHPs were captured at six sites: (a) Mombasa (4° 03′S, 39°40′E), a city at the coastal region of Kenya with a population of 1.2 million people. Mombasa has a tropical climate with hot and humid weather, temperature of 29°C and 11.2 mm of precipitation. (b) Kisumu (0° 00′N, 34°48′E), located in western Kenya at the shore of Lake Victoria has a population of about 599,468 people. The lake side town has no true dry season but significant rainfall throughout the year averaging 1321mm and temperatures of 24°C. (c) Murang’a (0° 43′ S, 37° 09′E) in central Kenya has warm and temperate climate (averaging temperatures of 17.4°C and 1590 mm of rainfall) and a population of about 1.06 million people. (d) NHPs were sampled from Kakamega County, Western Kenya. This region has a population of about 1.8 million people and temperatures average 20.4 °C with 1971mm of rainfall annually. Here, NHPs were captured from three peri-urban townships namely Buyangu (0° 19′N 34°57′E), Isecheno (0° 17′N 34°51′E) and Malava (0° 26′N 43°51′E).

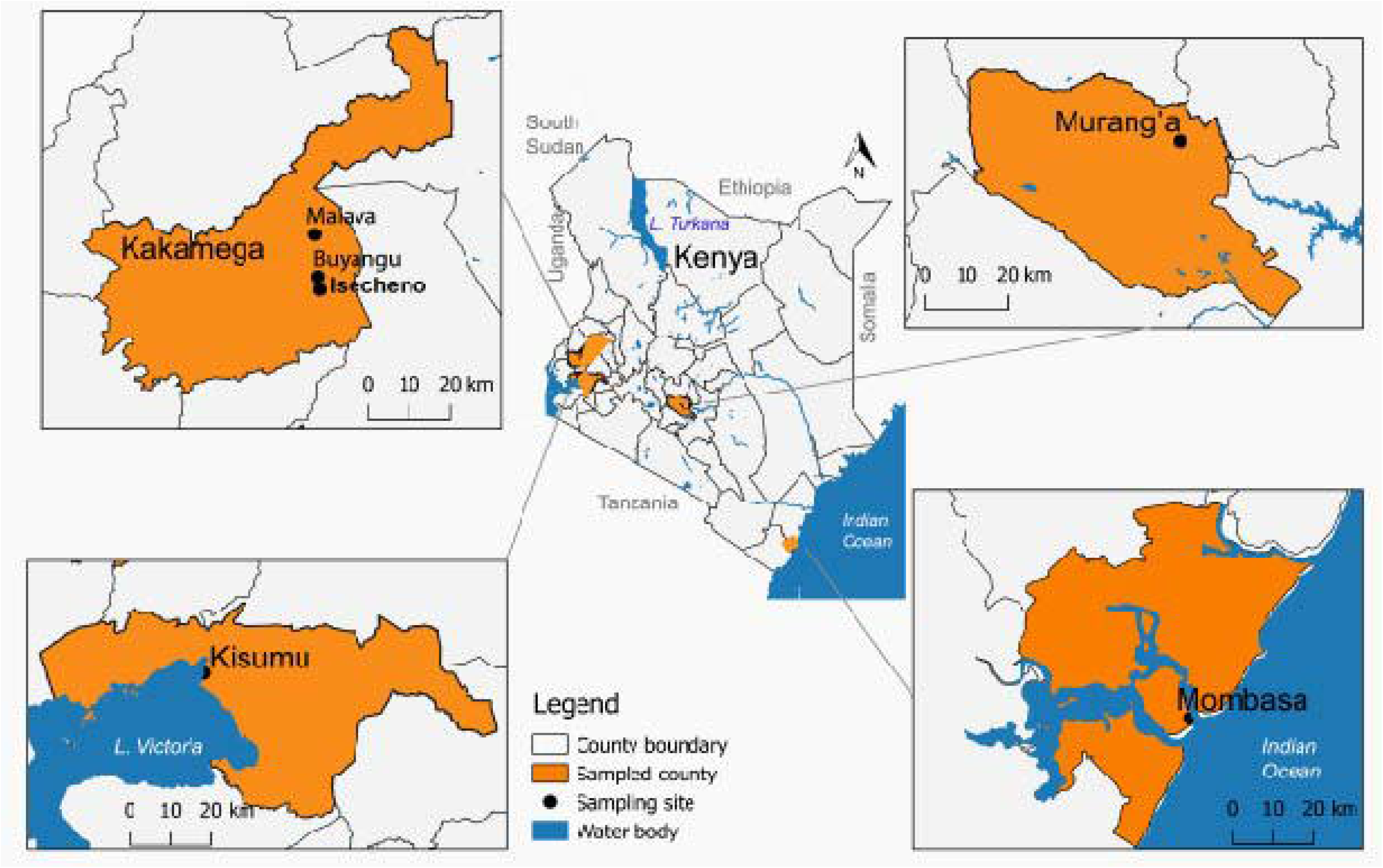

### Animal trapping and sample collection

Animal sampling was opportunistic. NHPs trapped were targeted for translocation to wildlife reserves because they were a public menace in urban areas or were regarded as pests by small-scale farmers within peri-urban areas. They were trapped as previously described (Maamun et al., 2011) following the guidelines for animal trapping and sampling after appropriate ethical review committee approval was received. Age group and sex of the trapped animals was determined according to Brett et al., (1982). Faecal samples were collected from the rectum via swabbing and stored in 70% ethanol for molecular assay. Where only one NHP was trapped in a cage, defecated material was collected and stored in 10% formalin for parasitological assays. If more than one animal was trapped in a cage, the faecal material in the cage was not collected because sample identity could not be confirmed. The samples were transported in a cool box with dry ice (−78.5°C) to the laboratory.

### NHPs sampled

A total of 86 NHPs: 41 African green monkeys (*Chlorocebus aethiops*), 30 olive baboons (*Papio anubis anubis*), 5 blue monkeys (*Cercopithecus mitis stuhlmanni*) and 10 red tailed monkeys (*Cercopithecus ascanius schmidts*) were trapped. The species caught from each location, their age category and sex are shown in Table 1. All 86 animals were rectal swabbed for samples while 69 that were individually trapped over the study period had extra faecal samples collected from their cages. In some cases where two or more animals were trapped in a single cage, it was impossible to utilise any of the excreta because of sample identity.

**Table 1:** The total number of NHP species sampled, distribution of each species according to sampling urban regions according to sex and age-group.

| Region | Sex | Age-group | AGM | Olive baboon | Blue monkey | Red tailed monkey |
| --- | --- | --- | --- | --- | --- | --- |
| Mombasa<br>30 | Male<br>18 | Juvenile | 1 |  |  |  |
|  |  | Sub/Adult | 9 |  |  |  |
|  |  | Adult | 8 |  |  |  |
|  | Female<br>12 | Juvenile | 2 |  |  |  |
|  |  | Sub/Adult | 2 | 1 |  |  |
|  |  | Adult | 7 |  |  |  |
| Kakamega<br>28 | Male<br>14 | Juvenile |  | 2 |  | 1 |
|  |  | Sub/Adult |  | 3 | 1 | 3 |
|  |  | Adult |  | 2 | 1 | 1 |
|  | Female<br>14 | Sub/Adult |  | 2 |  | 2 |
|  |  | Adult |  | 6 | 3 | 1 |
| Kisumu<br>21 | Male<br>10 | Juvenile |  | 1 |  |  |
|  |  | Sub/Adult | 2 | 3 |  |  |
|  |  | Adult | 1 | 3 |  |  |
|  | Female<br>11 | Sub/Adult |  | 1 |  |  |
|  |  | Adult | 2 | 6 |  | 2 |
| Murang'a<br>7 | Male<br>5 | Juvenile | 1 |  |  |  |
|  |  | Adult | 4 |  |  |  |
|  | Female<br>2 | Adult | 2 |  |  |  |
| 86 |  |  | 41 | 30 | 5 | 10 |

### Parasitological examination

Formal-ether sedimentation and sheathers sugar floatation techniques (Lee et al., 2010) were used for microscopic helminth detection. In both approaches, the slides were examined at 400X in duplicates in a Leica DM2000 LED microscope equipped with a digital camera control unit (Leica DFC 450) and representative images captured. Parasites were identified on the basis of egg color, shape, internal contents and larvae according to (Kouassi et al., 2015).

### DNA extraction, PCR assay and sequencing

A total of 86, ethanol-preserved, rectal swabbed faecal samples were snap-frozen in liquid nitrogen and ground to fine powder. The powder was homogenised with double distilled water as described previously (Phuphisut et al., 2016). Total DNA was extracted from 200 µl of the homogenate using the QIAamp DNA stool Mini kit (Qiagen, Hilden, Germany), according to the manufacturer’s instructions and stored at −20°C.

*Oesophagostomum* spp. were detected by PCR amplification of ITS2 gene using NC1 forward (5’-ACG TCT GGT TCA GGG TTG TT-3’) and NC2 reverse (5’-TTA GTT TCT TTT CCT CCG CT -3’) primer pairs (Ghai et al., 2014). PCR was performed in a RotorGene Q thermocycler (Qiagen, Hilden, Germany) using 10µM concentrations of each primer, 4µl of 5X HOT FIREPOL Eva Green HRM Mix (Solis Biodyne, Tartu, Estonia) and 2µl of the DNA template in a 25µl reaction mix. Thermal cycling conditions were as follows: initial denaturation at 95°C for 15 minutes, followed by 35 cycles of denaturation at 95°C for 1 min, annealing at 55°C for 45 secs, and extension at 72°C for 45 secs and a final extension at 72°C for 5 min. The PCR products were immediately utilised for high resolution melting (HRM) analysis as described (Villinger et al., 2016). Briefly, the amplicons were denatured at 95°C for 1 min, annealed at 40°C for 1 min and equilibrated at 65°C for 90 sec, and then increasing the temperature in 0.1°C increments up to 90°C with fluorescence acquisition after 2 seconds incremental holding periods. The melting curve profile was then analysed using Rotor-Gene Q series software version 2.1 with fluorescence (melting curve) normalised by selecting the linear region before and after melting transition. Melting temperature (Tm) was interpolated from the normalised data as the temperature at 50% fluorescence. Distinct HRM profiles, normalised in the range of 80-90°C, were visually determined for each reaction after completion of HRM data acquisition. PCR-HRM products were further visualised by 2% agarose gel electrophoresis stained with ethidium bromide. Gel readings were compared with corresponding PCR-HRM melting peaks for consistency with HRM analysis. Representative positive amplicons from PCR-HRM amplification were purified using the QIAquick Gel Extraction Kit (Qiagen, Hilden, Germany), according to the manufacturer’s instructions and sequenced.

### Phylogenetic analysis

Consensus sequences for ITS2 rDNA gene were generated from forward and reverse sequence data using Seqtrace version 0.9.0 (Stucky, 2012) and their identity ascertained by BLAST (Altschul et al., 1990) searches of GenBank (Benson et al., 2005). For species identification, a homology cut-off of 97-100% identity with a GenBank E-value threshold of 1e-130 was used. Sequences generated from this study and ITS2 ribosomal DNA sequences retrieved from GenBank were used for multiple sequence alignments in MAFFT (Edgar, 2004). Evolutionary analyses were performed to determine the relatedness and diversity of the *Oesophagostomum* spp. and *Trichostrongylus* spp. identified in this study to other nematode species endemic in East Africa. The evolutionary history was inferred using the maximum likelihood method in MEGA7 (Kumar et al., 2016) with bootstrapping at 1000 replicates. Phylogenetic trees were rendered in iTOL (Letunic & Bork, 2019).

## Results

### Microscopic nematode detection

Microscopic examination of the 69 faecal samples identified nematode eggs and/or larvae in 78.3% (54/69) of the samples. These included *Strongyloides* spp., *Ascarid* spp., *Trichuris* spp., *Oesophagostomum* spp., and *Enterobius* spp. (Figure 2A). *Strongyloides* eggs and larvae were observed in 25/69 (36.23%) of the samples while *Trichuris* eggs were observed in 54/69 (78.3%) of the samples. Eggs of *Enterobius, Ascarid*, and *Oesophagostomum* species. were each observed in 5/69 (7.2%) of the faecal samples (Figure 2B).

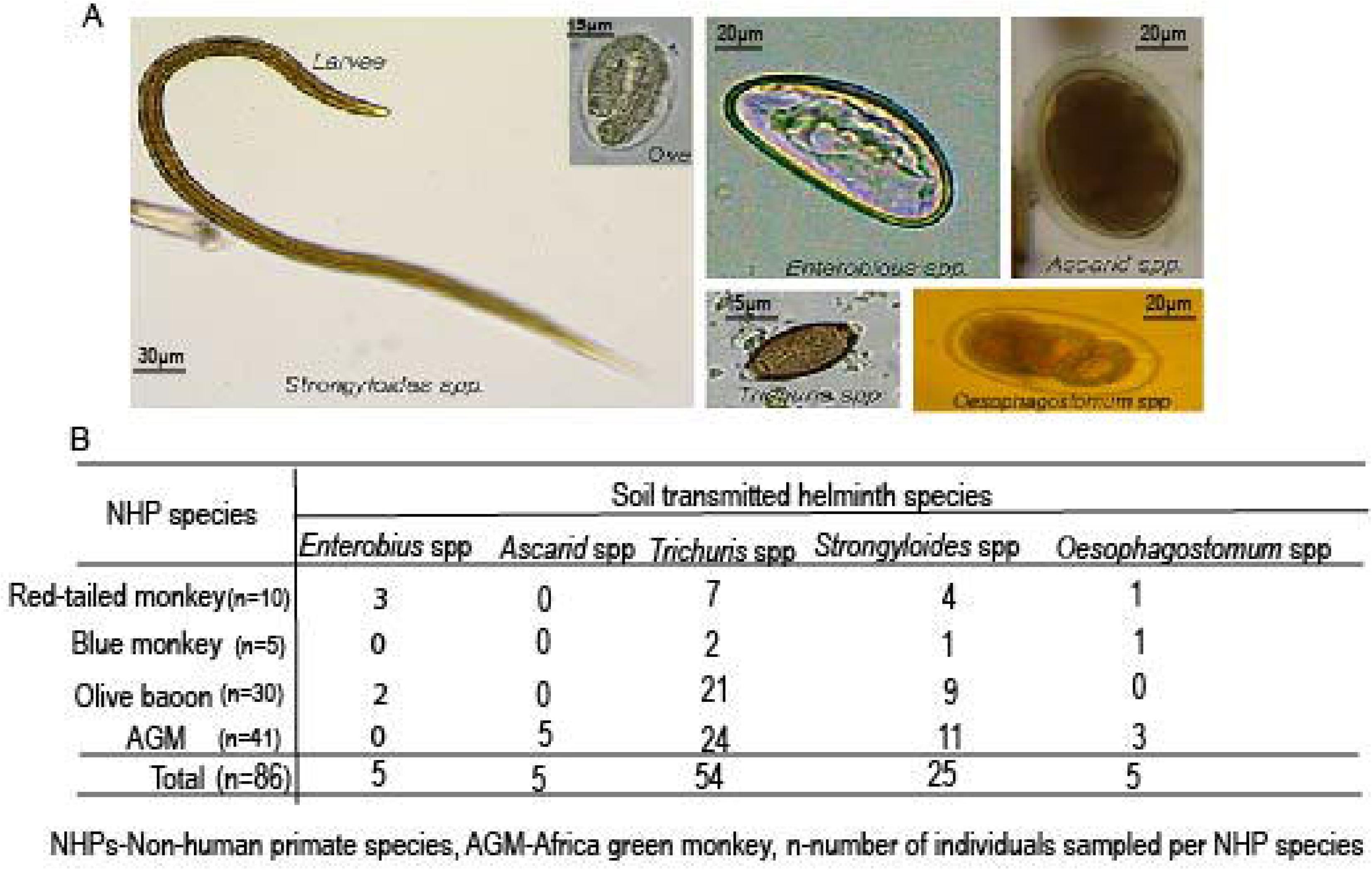

### Molecular detection of nematodes and sequencing

PCR-HRM analysis (Figure 3) identified *Oesophagostomum* spp. in 33/86 (38.4%) of the NHPs. Of these, 21 and 23 animals were infected with *O. bifurcum* and *O. stephanostomum* respectively while 11 animals were co-infected with both species (Table 2). BLAST searches of GenBank upon sequencing confirmed their presence and identified another nematode, *Trichostrongulus colubriformis*. These sequences have been submitted to the GenBank under accession numbers (MT184881 to MT184891).

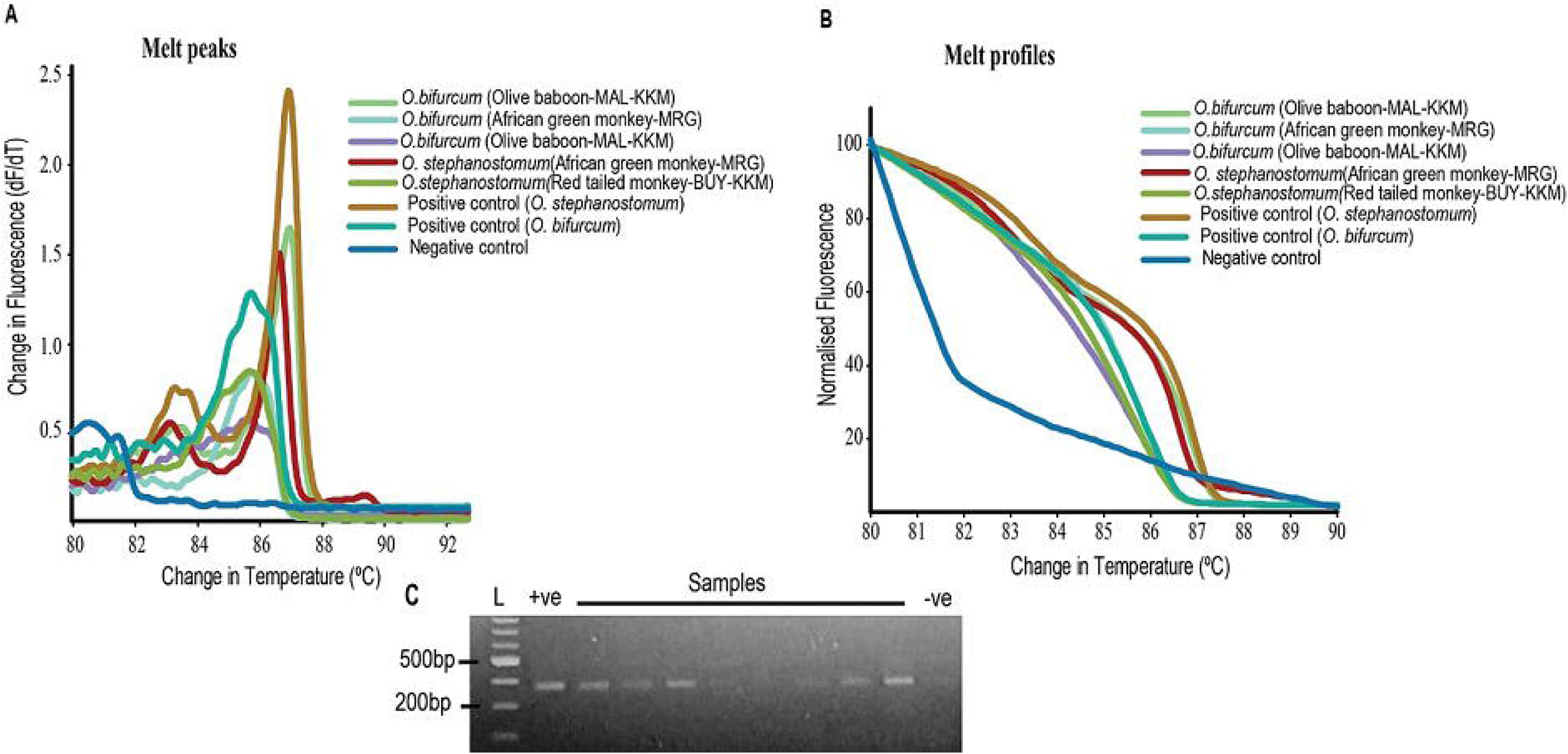

**Table 2:** *Oesophagostomum* spp. infection of non-human primates (NHPs) from urban and peri-urban centres in Kenya.

| Species | Number sampled | *O. stephanostomum* | *O. bifurcum* | *O. stephanostomum* and *O. bifurcum* |
| --- | --- | --- | --- | --- |
| Africa green monkey | 41 | 12(29.3%) | 9(21.95%) | 5(12.19%) |
| Olive baboon | 30 | 9(30.0%) | 4(13.33%) | 4(13.33%) |
| Blue Monkey | 5 | 1(20.0%) | 3(60.0%) | 1(20.0%) |
| Red-tailed monkey | 10 | 1(10.0%) | 3(30.0%) | 1(10.0%) |
| **Total** | 86 | 23(26.74%) | 21(24.42%) | 11(12.79%) |

### Overall nematode infection by microscopy and PCR-HRM

Parasitological and molecular assays enabled identification of nematodes in 74/86 (86.05%) NHPs samples. Overall, (35/41) AGMs, (27/30) olive baboons, (9/10) red-tailed monkeys and (3/5) blue monkey showed evidence of helminth infection. *T. colubriformis* and *Ascarid* spp. were only detected in AGMs, *Enterobius* in olive baboons and red-tailed monkeys while the rest occurred in all the NHPs (Figure 4). Two AGMs were infected with *T. colubriformis*. Infection varied across the sampling sites with 89.3% (25/28) in Kakamega, 81.0% (7/21) in Kisumu, 85.7% (6/7) in Murang’a and 86.6% (26/30) in Mombasa. *Ascarid* infections occurred only in Murang’a, *T. colubriformis* in Mombasa while *Enterobius* occurred in Kisumu and Kakamega.

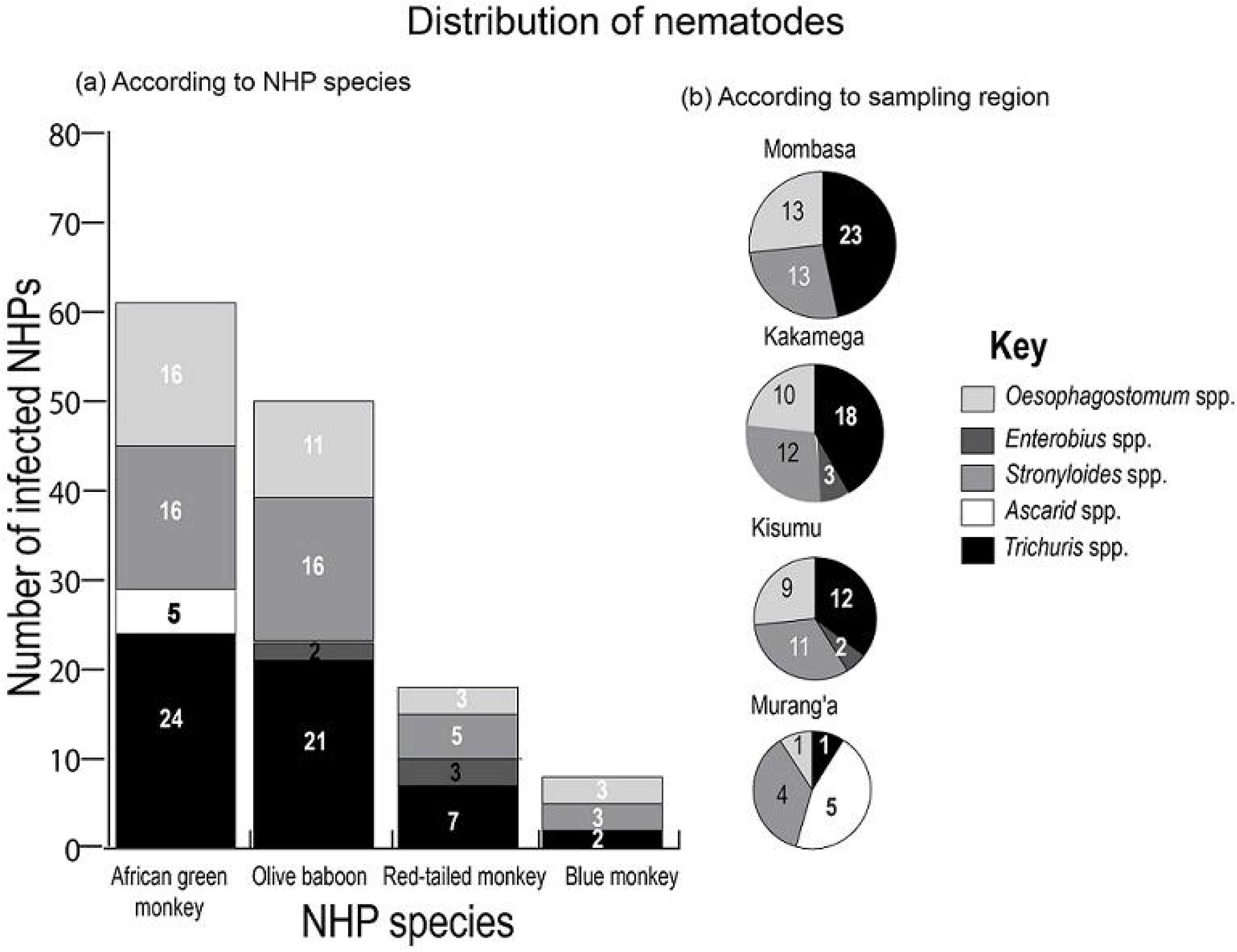

### Helminth co-infections as identified by microscopy and PCR-HRM

Co-infections were observed across the sampling sites in various NHPs (Figure 5). *Trichuris* and S*trongyloide*s. co-infections were the most common 17.44% (15/86) and occurred mostly in AGMs from Mombasa 24.39% (10/41). *Strongyloides* and *Oesophagostomum*. co-infection was also detected in 4.65% (4/86) of all the NHPs except AGMs. Co-infections with three nematodes were observed in all NHPs except the red-tailed monkey and were absent in NHPs trapped from Kisumu and Murang’a. Co-infections with four nematodes was observed in 8.12% (7/86) NHPs. Mixed infection with the two *Oesophagostomum* species occurred in 11 NHPs (Table 2).

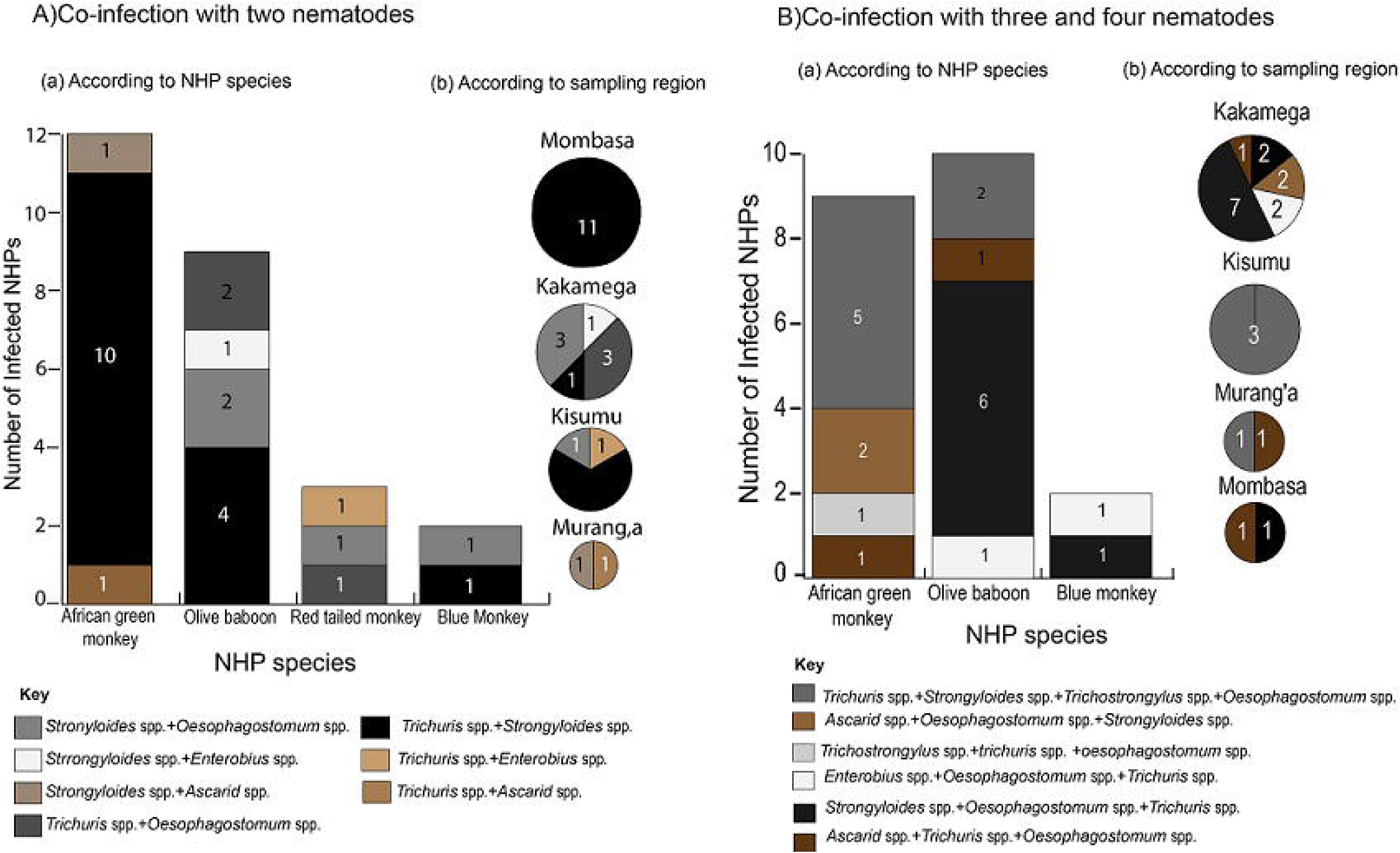

### Phylogenetic analysis

The phylogenetic tree obtained resolved into three distinct clades of *O. stephanostomum, O. bifurcum* and *T. colubriformis* (Figure 6). The *O. stephanostomum* clade lacked species or geographical sub-structuring; red-tailed monkeys’ sequences and a blue monkey sequence formed one sub-cluster while sequences from AGM and red-tailed monkey formed a second sub-cluster. Contrastingly, the *O. bifurcum* clade grouped the baboon and AGM isolates in different sub-clusters. The cryptic *Oesophagostomum* species described from Uganda (Ghai et al., 2014) was phylogenetically distinct from both *O. stephanostomum* and *O. bifurcum* identified in this study. The *T. colubriformis* clade grouped AGM-derived isolates in a distinct cluster from previous sequence of *T. colubriformis* DNA isolated from yellow baboon in Kenya and those reported from humans in Laos.

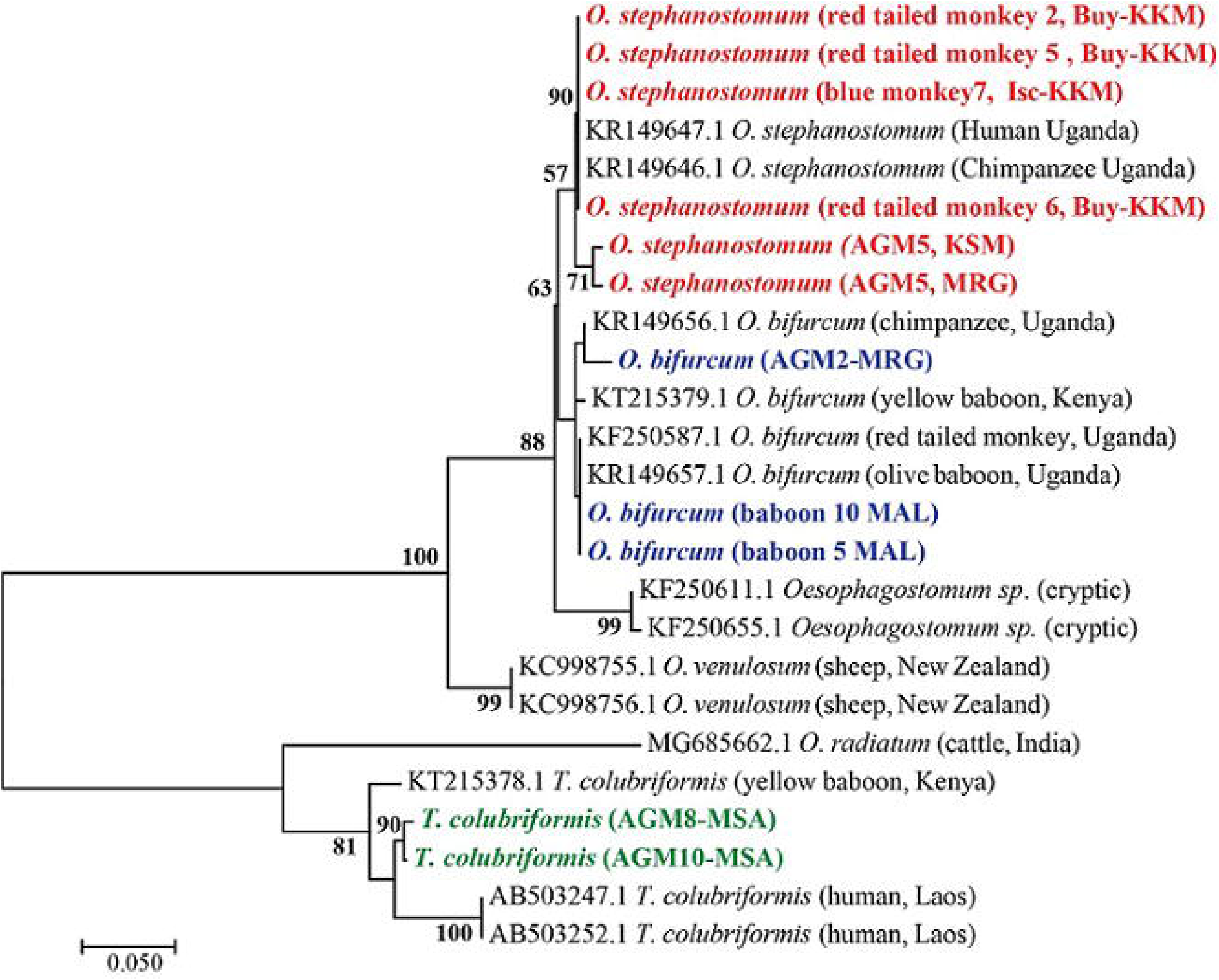

## Discussion

The diversity of STHs identified in selected Kenyan urban centres highlights the possible reservoir role played by NHPs in helminths urban ecology. The significant diversity of nematodes infecting different species of NHPs within urban and peri-urban centres compliments previous molecular surveys that detected helminths in free-ranging NHPs within remote wildlife habitats in Kenya (Akinyi *et al*., 2019; Obanda *et al*., 2019). In STHs endemic countries, re-infection has been observed in half of the children treated for intestinal worms (Jia et al., 2012) a phenomenon attributed to re-infection due to persistence of infective worm stages in the environment. As the Kenyan government commits to strengthen helminth control (Mwandawiro et al., 2019) it is faced with the challenge of identifying reservoir hosts. This study provides useful information on possible reservoir hosts of STHs that may contribute to environmental sustenance of zoonotic nematodes in urban and periurban regions of Kenya. It also illustrates the possibility of utilising PCR-HRM analysis to efficiently differentiate between *Oesophagostomum* spp. without the need for sequencing as earlier described for hookworms (Ngui et al., 2012). PCR-HRM differentiated *O. stephanostomum* and *O. bifurcum* co-infecting NHPs (Figure 3) as confirmed by sequencing indicating its utility as a non-subjective approach to supplement sequencing for accurate characterisation of nodular worms.

Phylogenetic reconstruction of nodular worm isolates demonstrated separation of *O. stephanostomum* sequences into a sub-cluster consisting of isolates from red-tailed monkeys and blue monkeys in Kakamega and a sub-cluster consisting of sequences from AGMs sampled in Kisumu and Murang’a indicating that *O. stephanostomum* is neither geographical nor host species-based. In addition, clustering of *O. stephanostomum* detected in this study with an isolate from human (accession number KR149647.1) indicates close evolutionary relatedness and therefore suggests the potential for this parasite to infect both humans and NHPs. This finding concurs with Cibot et al. (2015) reports of human and NHP infection with *O. stephanostomum*, suggesting increased risk of transmission for this helminth between primate species. *O. bifurcum* sequences from olive baboons in this study formed a monophyletic cluster with *O. bifircum* sequences from other NHP species retrieved from Genbank. However, the *O. bifurcum* sequence from a chimpanzee sampled in Uganda clustered with sequences recovered from an AGM in Kenya. The two were evolutionarily divergent from the rest of the sequences within this cluster. This points to potentially new host species since *O*.*bifurcum* has not been previously reported in AGMs in Kenya and is not commonly described in chimpanzees. Because transmission occurs via the ingestion of the infective third-stage larvae present in contaminated food or water, oesophagostomosis is a potential zoonotic risk when infected NHPs and humans share the same habitats. Therefore, intervention strategies to combat oesophagostomosis need to factor NHPs as potential reservoirs. Sequencing data also confirmed infection by *T. colubriformis* in olive baboons. *T. colubriformis* sequences generated in this study were distinct from the *Trichostrongylus* spp. isolates from yellow baboon in Kenya (Obanda et al., 2019)which may be as a result of sub structuring according to host species.

Other nematodes identified in this study included *Trichuris*., *Enterobius*., *Ascarid*., *Strongyloides*. and *Trichostrongylus* species. *Trichuris trichiura*, has been detected in different primate species in Kenya (Mbora & McPeek, 2009; Akinyi et al., 2019) and experimentally transmitted from NHPs to humans (Monteiro et al., 2007) providing evidence of its zoonotic potential. In addition, a single taxon was found to be both human and NHPs infective in a *Trichuris* host diversity study (Ghai et al., 2014). NHPs are known to be the major hosts of *S. stercoralis* and, especially, *S. fuelleborni*. A peculiarity of *Strongyloides* spp., is their ability to penetrate the host’s skin and/or autoinfect making them burdensome helminths causing long-term suffering. Identification of *Strongyloides* in this study is thus of public health concern. Klaus et al.,(2017) demonstrated *Ascarid* spp. cross-infection and zoonotic potential between human and NHPs. Nematodes in the genus *Trichostrongylus* are known to infect humans, wild animals, herbivorous animals (Obanda et al., 2019). Primates including humans become infected due to environmental contamination. Where NHPs, ruminants and humans live in sympatry, *T. colubriformis* is a perpetual public health burden thus its presence poses a potential risk of transmission in the study areas. Contrary to previous studies (Munene et al., 1998; Obanda et al., 2019) AGMs in this study were infected with *O. bifurcum* which adds AGMs as reservoir host species to the epidemiology of nodular worm. Overall, a complex anthropo-zoonotic transmission cycle may be maintained in the study regions.

Consistent with other studies (Klaus et al., 2017; Akinyi et al., 2019), we also observed coinfection with multiple nematode species. Parasites co-occurring within a single host interact in a variety of ways that influence their abundance, distribution, and the dynamics of one another (Pedersen & Fenton, 2007). One such interaction is the immune modulation. A helminth induced immunosuppression caused by infection with one helminth species may strongly alter a host’s response to subsequent infection by other species enhancing the likelihood of coinfection (Cox, 2001). Further, mechanisms related to host exploitation can be modified to ultimately enhance coexistence. For instance, the distinct spatial niches of worms within a host e.g. *Ascaris lumbricoides* predilection in jejunum while *Trichuris trichiura* resides in the cecum reduces levels of resources competition between the two worms. Predominance of *Trichuris* and *Strongyloides* co-infections in this study may be due to their abundance in the faecal samples as they were the most dominant infections. Given the ubiquity of coinfection in nature, and the effects coinfecting parasites are likely to have on one another, interactions among parasites may be a major force generating variation in the transmission of disease and in shaping infectious disease dynamics. While most studies on intestinal helminths of free-ranging NHP rely on faecal samples collected after defecation in the wild, challenges in interpretation of results may arise due to collection of several samples from the same animal (Gillespie et al., 2010). In the current study, stool samples could be traced back to the individual animal having collected the sample from the rectum of the animals or freshly excreted faecal sample from trapping cages with single animal.

With respect to host species infections, the taxonomic diversity was highest in AGMs followed by olive baboons, red-tailed monkeys and blue monkeys. This may be due to behavioral factors; blue monkeys are mainly arboreal while AGMs are semi-terrestrial, which means they spend more time on the ground hence increasing their chances for contact and contraction of infective stages of different nematodes. The rich taxonomic diversity in AGMs parallel other studies that report high prevalence of helminth infections in terrestrial NHPs (Poulin & Morand, 2005;Ghai et al., 2014; McCabe et al., 2014). Although olive baboons are also terrestrial, the slightly lower number of different nematode taxa identified compared to AGMs could be the consequence of sample size, and therefore strong inferences cannot be made. However, and to the best of our knowledge, studies comparing taxonomic diversity and infection rates among different species of NHPs in Kenya remain scarce.

The number of nematode infections in NHPs was highest in Mombasa and Kakamega while Murang’a had the least cases of infections. By comparison, the prevalence of all STHs in human population in Kenya is highest in coastal and western regions (Brooker and Michael, 2000; Masaku *et al*., 2017). One explanation for high cases of infection in Mombasa could be a coastal habitat providing ideal environmental conditions such as humidity and warmth for egg development. Although Kisumu is also warm, humidity is much lower. Additionally, frequent recurrence of STHs at the same location may facilitate environmental accumulation of infective parasite stages and could result in reinfection. This is especially relevant in the case of reservoir hosts, since adult worms’ lifespan is typically longer than annual periods when environmental conditions favours transmission thus maintaining overall endemicity.

A key limitation of this study is that we were unable to sample humans within the studied regions. Our sampling was opportunistic because we focused on NHPs that had been targeted for translocation to national reserves. In most cases, surveillance of zoonotic pathogens under the ‘One Health’ approach recommends that humans, wildlife and domestic animals that share same habitats are screened concurrently. Our approach to phylogenetic analysis addressed this limitation by comparing our generated sequences to those of nematodes isolated from humans and domestic animal faecal material. Although our data provides baseline information on the potential zoonotic risk of gastrointestinal nematodes in urban centres, we recommend further surveillance of the nematodes using the one health approach. Since NHPs serve as sentinels for surveillance of emerging diseases, the rich taxonomic diversity of nematodes detected NHPs from selected urban centres in Kenya could be an important reference to the helminth prevalence in altered urban habitats. In addition, the detection of *Oesophagostomum* species in free -ranging NHPs within densely populated urban centres is of public health interest because of the zoonotic nature of these nematodes, especially *O. bifurcum*, a parasite that can be lethal to humans.

## Authors Contribution

MJ, VO, PM, MA formulated the project, MJ, MA, IL, PM GE, conducted fieldwork, PM, MJ, MA, conducted laboratory analysis, MJ, EM, VO, RN, PM analysed the data. MJ, MA, PM, VO, EM drafted the manuscript. All the authors revised the manuscript and approved the final manuscript draft.

## Acknowledgements

The authors gratefully acknowledge the institutional support of the Institute of Primate Research (IPR), Kenya and the Kenya Wildlife Service (KWS). We also thank Dr. Moses Otiende and Mr. Samson Mutura for their assistance with the molecular experiments. This study received financial support from the Consortium for National Health Research (CNHR) grant, Kenya (Project number RCDG-041/2012) and by a Global Enhancement Fund grant, USA (2017) held by Prof. Susan Alberts, Duke University.

## Ethical Note

Animal trapping, sedation and sampling were undertaken with approval from and according to the guidelines of Institute of Primate Research (IPR) Institutional Scientific Review Committee and the Department of Veterinary and Capture Services of the Kenya Wildlife Service (KWS), Nairobi (Review number, ISERC/04/18).

